# Preictal dysfunctions of inhibitory interneurons paradoxically lead to their rebound hyperactivity and to Low-Voltage-Fast onset seizures in Dravet syndrome

**DOI:** 10.1101/2023.09.20.558646

**Authors:** Fabrizio Capitano, Mathieu Kuchenbuch, Jennifer Lavigne, Rima Nabbout, Massimo Mantegazza

## Abstract

Epilepsies have numerous specific mechanisms. Understanding neural dynamics leading to seizures is important for disclosing pathological mechanisms and developing targeted therapeutic approaches. We investigated electrographic activities and neural dynamics leading to convulsive seizures in patients and mouse models of Dravet syndrome (DS), a developmental and epileptic encephalopathy in which hypoexcitability of GABAergic neurons is considered to be the main dysfunction.

We analyzed EEGs from DS patients carrying a *SCN1A* pathogenic variant, as well as epidural electrocorticograms, hippocampal local field potentials and hippocampal single-unit neuronal activities in *Scn1a*^+/-^ knock-out and *Scn1a*^RH/+^ knock-in DS mice.

Strikingly, most seizures had low-voltage-fast onset in both patients and mice, which is thought to be generated by hyperactivity of GABAergic interneurons, the opposite of the main pathological mechanism of DS. Analyzing single unit recordings, we observed that temporal disorganization of the firing of putative interneurons in the period immediately before the seizure (preictal period) precedes the increase of their activity at seizure onset, together with the entire neuronal network. Moreover, we found early signatures of the preictal period in the spectral features of hippocampal and cortical field potential of *Scn1a* mice and of patients’ EEG, which are consistent with the dysfunctions that we observed in single neurons.

Therefore, the perturbed preictal activity of interneurons leads to their hyperactivity at the onset of generalized seizures, which have low-voltage-fast features that are similar to those observed in other epilepsies and are triggered by hyperactivity of GABAergic neurons. Spectral features may be used as predictive seizure biomarker.

**Significance statement:** Dravet syndrome (DS) is caused by mutations of the Na_V_1.1 sodium channel (*SCN1A* gene) leading to hypoexcitability of GABAergic interneurons. We found that most of the seizures in both DS patients and mouse models have low-voltage-fast onset, which is instead thought to be generated by hyperactivity of GABAergic neurons. We disclosed a disorganization in the temporal pattern of the firing of single interneurons before the seizure (preictal period), and a rebound hyperactivity at seizure onset. Consistently, the electrographic signal showed a decrease of fast oscillations in the preictal period. Thus, perturbed interneurons’ preictal activity, consistent with the main mechanism of DS, leads to their hyperactivity at seizure onset and induces specific electrographic signatures that may be exploited for seizure prediction.

## Introduction

Understanding neural dynamics leading to seizures is important for disclosing pathological mechanisms of epilepsy and for developing new therapeutic approaches. Epilepsy etiology is diverse with numerous underlying specific mechanisms. However, it is not yet clear if the same dynamics of neuronal subpopulations is shared in different epilepsies or if the correlation between the electrographic properties of seizures and the underlying neuronal dynamics is consistent between different models. Shedding light on these features can be particularly important for developmental and epileptic encephalopathies (DEE), in which seizures are often drug resistant and may worsen cognitive and behavioral outcomes (1, 2).

Dravet syndrome (DS) is a DEE characterized by onset in the first year of life with seizures triggered by fever/hyperthermia, which persist also in adulthood, and later appearance of afebrile seizures, as well as of developmental plateauing with behavioral disorders(3–6). *SCN1A* pathogenic variants leading to loss of function of the Na_V_1.1 sodium channel are identified in most individuals with DS(4, 7–10). Gene targeted mouse models replicate the phenotype of the patients and have been instrumental for showing that loss of function of Na_V_1.1 leads to hypoexcitability of GABAergic neurons, reduced inhibition and hyperexcitability of neuronal circuits, which is the main DS pathogenic mechanism thus far identified(8, 11–14). Notably, investigations in individuals with DS have shown reduced GABAergic inhibition *in vivo* (15). DS fits in the group of epilepsies characterized by both generalized and focal seizure(2). Indeed, DS patients show multiple types of difficult-to-treat seizures that are most frequent in infancy and childhood, including generalized (clonic, tonic-clonic, atypical absences, tonic and myoclonic) and focal (unilateral clonic, focal seizure with or without generalization, and myoclonic) seizures (5, 6, 16). As in other DEEs, in addition to their impact per se, seizures can contribute to phenotype worsening in *SCN1A* diseases (1, 2). Interestingly, in knock-in *Scn1a*^RH/+^ mice, an asymptomatic phenotype is transformed into a DS-like one upon induction of few short seizures(17).

Little is known about neuronal dynamics in DS seizures. Previous studies in both patients and models of other types of epilepsies have identified specific seizure onset patterns and investigated the involvement of different neuronal subpopulations(18–22). In particular, the analysis of electrographic recordings of focal seizures has led to the identification of two main onset patters. Low-voltage fast-onset (LVF) discharges initiate with 1 or 2 interictal-like population spikes (denominated “sentinel spikes”) followed by low-amplitude, high-frequency activity; hypersynchronous-onset (HYP) discharges initiate instead with a longer series of population spikes in the period immediately preceding the seizure (preictal period) without low-amplitude, high-frequency activity(18–22). Optogenetic studies showed that activation of GABAergic neurons can directly cause LVF seizures, whereas activation of excitatory pyramidal neurons can drive HYP seizures(23, 24).

We have analyzed here the electrographic activity recorded during generalized convulsive seizures (GCS) in both DS patients and gene targeted mouse models. We surprisingly observed seizures with LVF onset. In addition, we studied the neuronal dynamics that lead to this type of seizures in DS and observed specific perturbations of the activity of putative inhibitory interneurons in the preictal period, which precede the increase of their activity at seizure onset.

## Results

### DS mice and patients show convulsive seizures with Low Voltage Fast (LVF) onset

We recorded, during spontaneous or hyperthermia-induced convulsive seizures of *Scn1a*^+/-^ mice, the local field potential (LFP) in the dorsal hippocampus, which is implicated in seizure generation in DS mouse models(25–27). Surprisingly, most onsets of both spontaneous and hyperthermia-induced seizures could be classified as Low Voltage Fast (LVF) by the presence of the “sentinel” spike heralding the typical fast oscillations of low amplitude around the onset (Fig.1A-B-E). In particular, 44 out of 53 spontaneous seizures (6 mice) and 24 out of 31 hyperthermia-induced seizures (3 mice) were with LVF onset. The remaining seizures were difficult to classify according to established onset subtypes(28, 29). Since the signal recorded in patients by scalp EEG mainly provides a measure of neocortical activity, we investigated whether the LVF onset in Scn1a^+/-^ mice is a specific feature of the hippocampal signal or could be a generalized property of cortical networks. Thus, we analyzed the electrocorticogram (ECoG) recorded from the parietal cortex in a separate group of Scn1a^+/-^ mice during spontaneous seizures. Also in this case, most of the seizures (52 out of 66 in 9 mice) showed LVF onset, confirming that this is a general property of seizures in DS mice (Fig.1C). We found no significant differences in the proportion of LVF onsets observed in the three experimental conditions (Fisher exact test p=0.57; Fig.1D).

**Figure 1.**
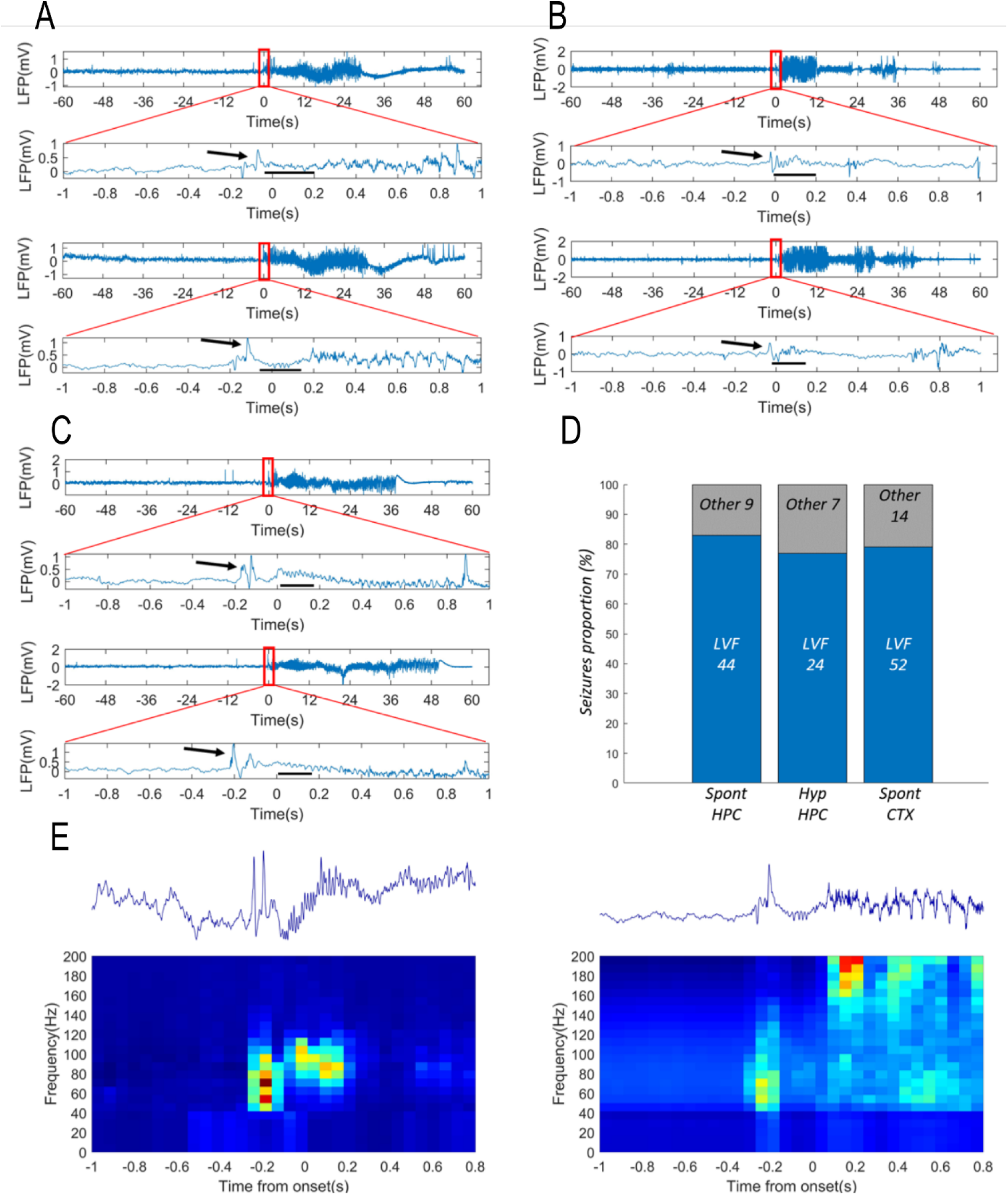
Spontaneous and hyperthermia-induced seizures have LVF onset in Dravet mice. Representative traces of hippocampus LFP recorded during spontaneous seizures (**A**) or hyperthermic seizures (**B**) with LVF onset recorded in Scn1a^+/-^ mice. The black arrows indicate the “sentinel” spike. The black dashes underline the Low Voltage Fast (LVF) oscillations. The signal around the onset was manually checked and the specific onset point placed at the beginning of abnormal activities. In the case of low-voltage fast pattern, representing the majority of the observed onsets, this was placed at the beginning of LVF activity. **C**. Electrocorticogram (ECoG) recorded on the parietal cortex of Scn1a^+/-^ mice during spontaneous seizures, which showed properties that were similar to those of hippocampal LFP. **D.** Bar-chart plot displaying the proportion of seizures with LVF onset in the LFP signal recorded in the hippocampus during spontaneous seizures (Spont HPC) or hyperthermia induced seizures (Hyp HPC), and in the ECoG signal during spontaneous seizures (Spont CTX). **E.** Two examples of LFP spectrogram of LVF onset seizures showing the low-voltage fast activity localized in the gamma band.

We then analyzed the onset features of 11 GCS recorded in 7 DS patients with *SCN1A* pathogenic variants. The mean individuals’ age was 8.4 years old [range 5.5-10 years]. Clinically, seizures were generalized tonic vibratory (n=6), generalized tonic with secondary head and eyes deviation (2), and generalized tonic-clonic (2). Similar to the mouse model, also in patients we identified in 54.5% of seizures the prototypical features of the LVF onset (Fig.2).

**Figure 2.**
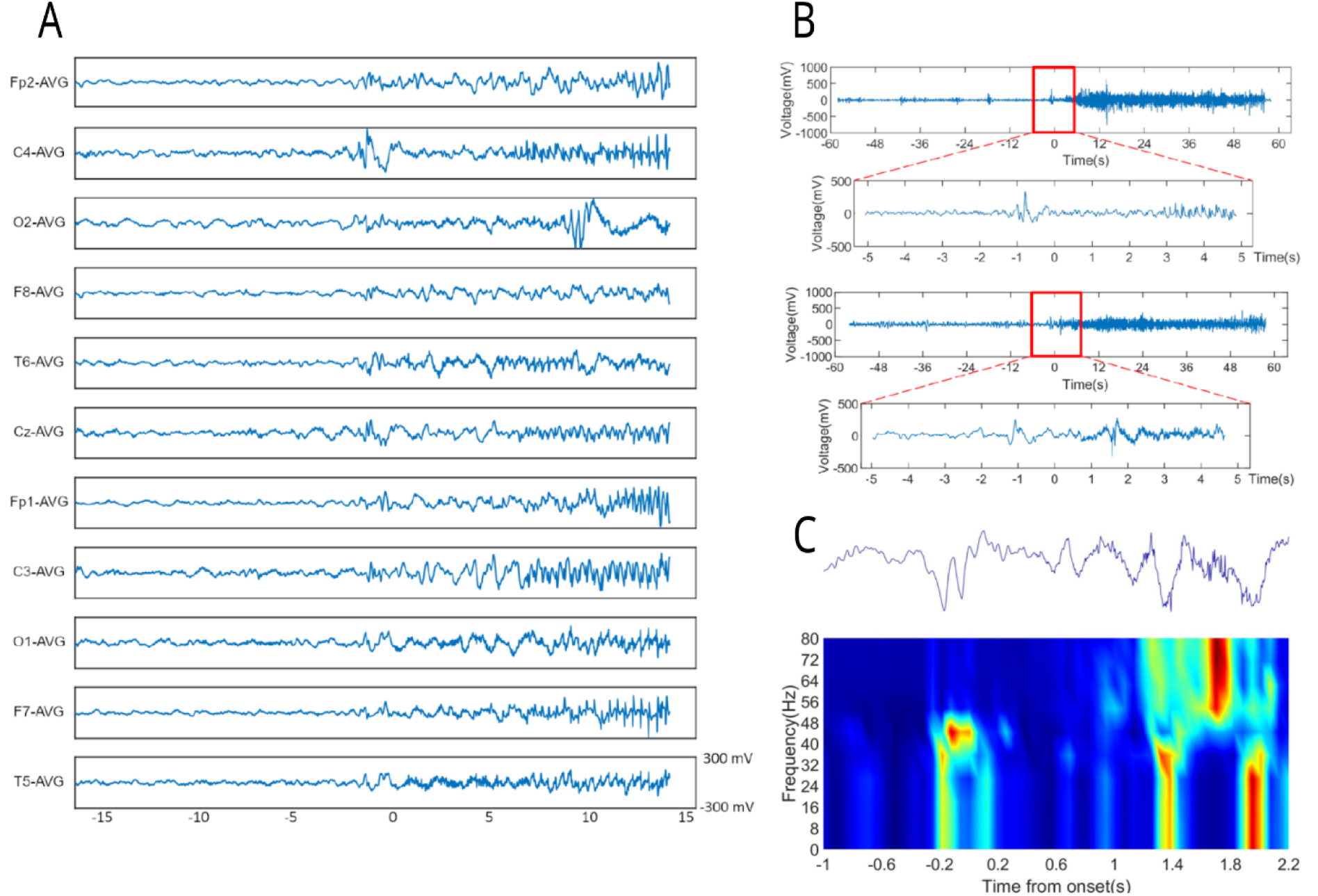
The onset of most of the seizures in individuals with Dravet syndrome in our cohort showed LVF onset, as in mice. **A**. Example of seizure onset on the 11-channel EEG of a 10.8-year-old boy with Dravet syndrome and LVF seizure onset pattern (a sentinel spike followed by a low-voltage fast activity) (individual n°3). **B**. Upper panel; single-channel EEG (C4) illustrating the onset of a LVF seizure in a 5.5-year-old boy with Dravet syndrome (individual n°2). Lower panel; single-channel EEG (C4) illustrating the onset of the LVF seizure in a 9.1 –year-old boy with Dravet syndrome (individual n°6). A zoom on the 12s around the onset of the seizure (red rectangle) is provided for both traces. The black arrows indicate the “sentinel” spike; the black segments highlight the Low Voltage Fast (LVF) oscillations. **C.** Example of LFP spectrogram of LVF onset seizures showing the low-voltage fast activity that is localized in the gamma band.

### Seizures in DS mice recruit different neuronal populations

These results are unexpected and puzzling because the proposed main pathological mechanism of DS, GABAergic neurons’ hypoexcitability, should hinder the ictogenic mechanism of LVF onset seizures, which has been linked to GABAergic neurons’ hyper-activity(23); on the other end, preponderant HYP onsets would be expected in case of hyperactivity of glutamatergic neurons(18–22).

To shed light on this issue, we recorded *in vivo* the activity of single neuronal units with tetrodes in the dorsal hippocampus of *Scn1a*^+/-^ mice during both hyperthermia-induced and spontaneous seizures. We identified by spike-sorting 142 single neurons and we classified them based on their spike duration to investigate the involvement of putative inhibitory and excitatory neurons in driving seizures onset. The distribution of action potential durations displays a bimodal shape (Fig.3A), consistently with previous studies (30–33). Thus, we classified sharply spiking neurons as putative inhibitory neurons and widely spiking neurons as putative excitatory neurons using 400ms as a threshold. Notably, this threshold is coherent with multiple previous observations (30–33). To cross-validate the classification we demonstrated that putative inhibitory neurons have a shorter hyperpolarizing phase, as measured by AHP half-width, as well as a higher average firing frequency (Fig.3A-B; Mann-Whitney test for AHP, U=1641, p<0.0001; for average frequency U=6242, p<0.0001).

**Figure 3.**
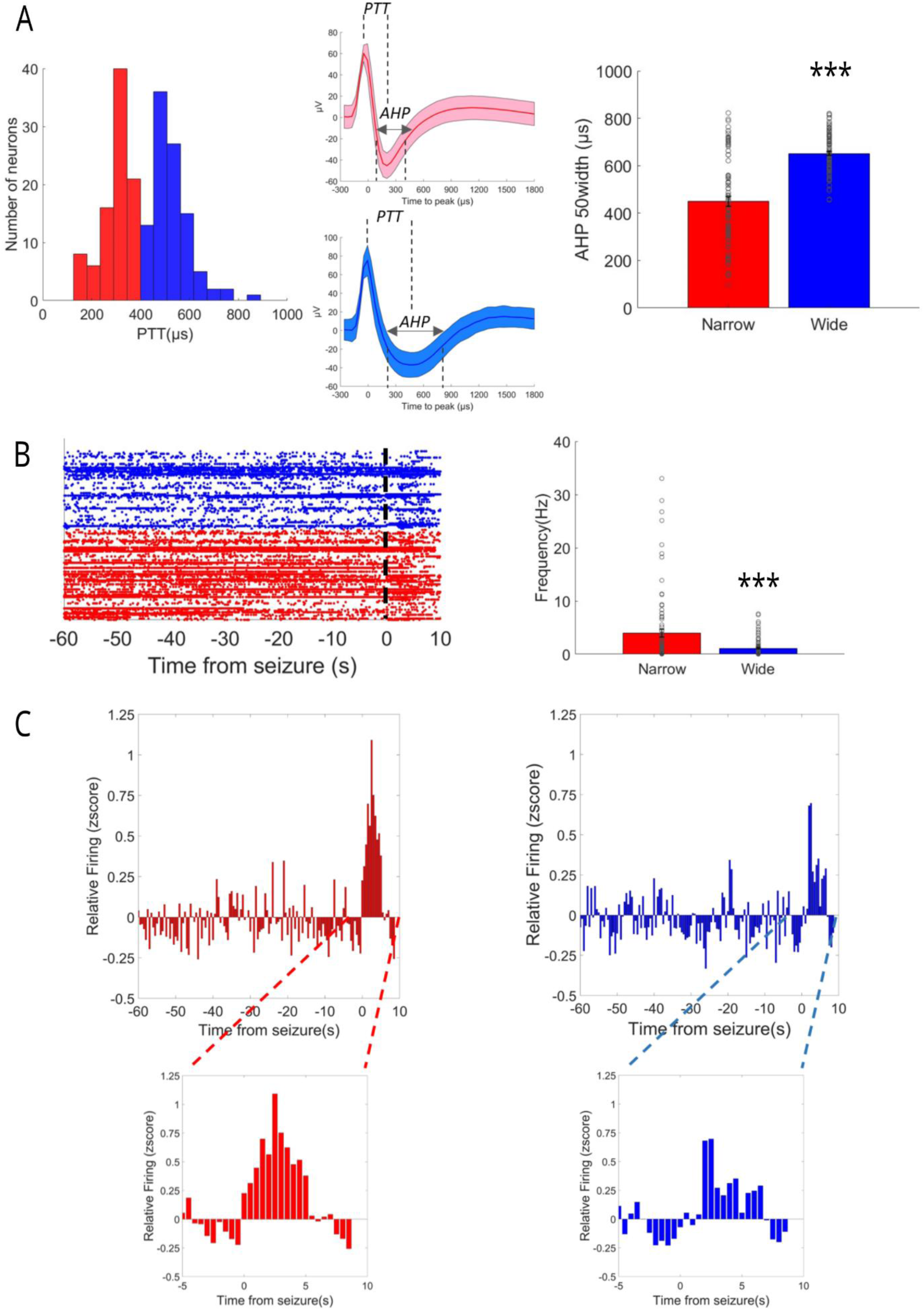
Dynamics of single neuronal units in LVF seizures of Dravet mice. **A.** Identification of putative neuronal subpopulations by action potential shape. The spike duration was evaluated by peak-to-trough duration (PTT) and shows a bimodal shape, consistent with the presence of sharp spiking (red) and widely spiking (blue) neurons. In the middle, two examples of sharp spiking and widely spiking neurons are showed. The line represents the average action potential shape, the shaded area the standard deviation. The analysis of the after-hyperpolarization (AHP) duration and of the average firing frequency revealed a significant difference between the two populations **B.** Representation of the neuronal activity of the two putative neuronal populations. The analysis of average firing frequency revealed an higher activity of sharply spiking neurons. **C**. Quantification of neuronal activity in the preictal period and at onset of seizures induced by hyperthermia. The inserts on the bottom show for each subpopulation the activity zoomed around the seizure’s onset. ***, p<0.001

The quantification of the preictal firing frequency showed that both populations increase their average activity at seizures onset (Fig.3C; Mixed model ANOVA for sharply spiking neurons F137,9727=2.92, p-value<0.0001; for broadly spiking neurons F137,9453=1.72, p-value<0.0001). Given that the averaging hides inter-neuronal variability, we quantified the proportion of neurons sustaining seizures (defined as the neurons that displayed activity with a z-score>1.96 for at least two consecutive 500ms bins during the seizure). The two neuronal subpopulations showed no statistical difference: 26 out of 72 putative inhibitory neurons and 16 out of 70 putative excitatory neurons were classified as active after the seizure onset (Fisher exact test p=0.1). The proportion of active neurons of the two subpopulations was not statistically different when spontaneous and hyperthermia-induced seizures were compared (Fisher exact test p=0.26 and p=0.16 for respectively sharply spiking and widely spiking neurons) suggesting that neural mechanisms at the seizure’s onset are shared between the two types of seizures.

Previous studies suggested that even when LVF seizures are initiated by hyper-activity of interneurons, principal cells can subsequently sustain the ictal activity during the seizure (18). Even if this temporal shift in our recordings is not evident on the average activity of the two subpopulations at the seizure onset (Fig.3C), we further investigated this issue analyzing individual neurons. More specifically, for cells classified as “active” during the seizure, we identified the first active time bin after the seizure’s onset and we compared sharply spiking and widely spiking neurons. We didn’t find any differences in the time of recruitment between them (t-test p-value p=0.8), confirming the observation on the average population.

### Seizures in DS mice are preceded by early perturbations of EEG, LFP and putative inhibitory interneurons’ firing

The activity of the network during seizures is informative about the circuits actively sustaining the ictal discharges once they are triggered, but it is not necessarily correlated to the perturbations occurring before seizure’s onset, which lead the network toward the ictal state. Thus, we analyzed LFP and single neuronal activity in the preictal period preceding the spontaneous seizures.

We first focused on the analysis of the hippocampal LFP and of the parietal cortex ECoG of DS mice, performing spectral analysis to quantify the power of different frequency bands and its dynamics in the preictal period (see methods for details). We found that spectral properties of both hippocampal (Fig.4A) and cortical (Fig.4B) neural signal are modified in the minutes preceding the seizures. In particular, we observed an early and progressive increase of slow rhythms and theta oscillations and a decrease of fast oscillations, which anticipate the increase of the total signal power occurring a few seconds before seizure onset.

**Figure 4.**
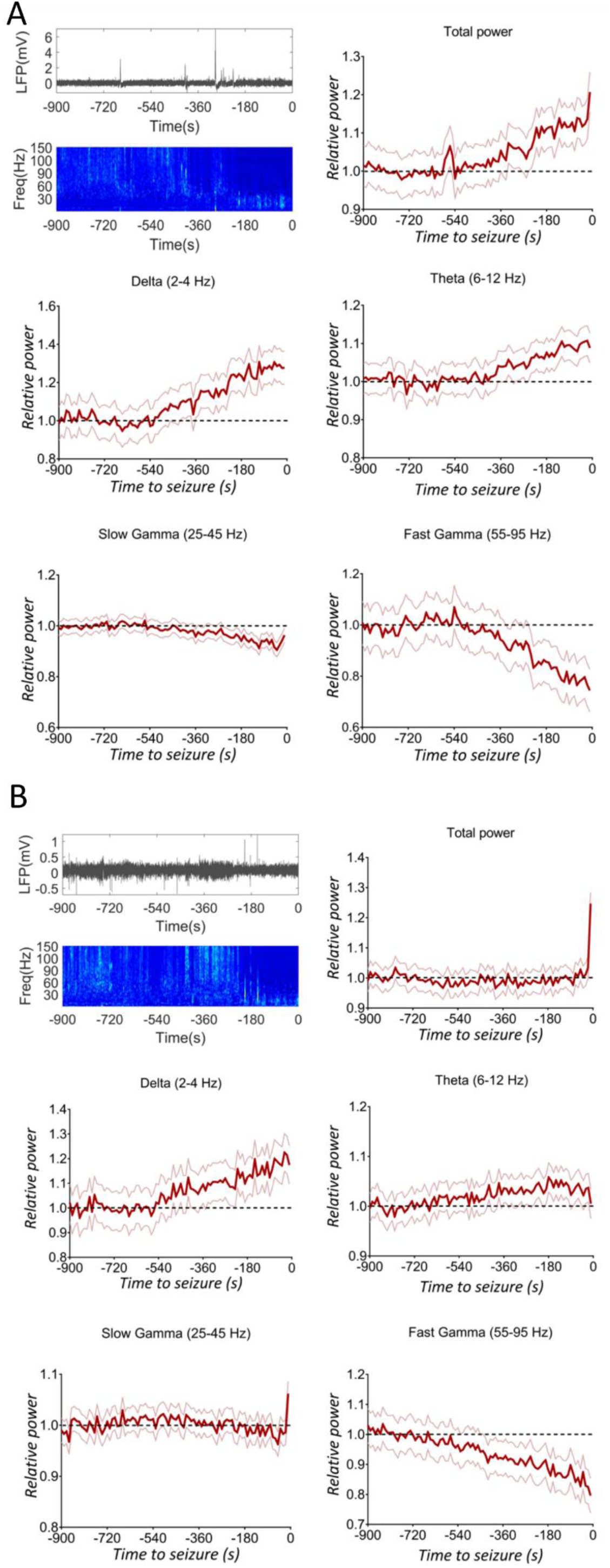
Seizures are preceded by early modifications of LFP spectral properties in mice. **A**. Representative hippocampal LFP recording in the preictal period with its corresponding spectrogram and quantification of mean spectral power decomposed in different frequency bands (the 95% confidence interval for the different time points is displayed), showing an increase for slow rhythms and theta oscillations and a decrease for fast oscillations (mixed model repeated measures ANOVA, p<0.0001 for delta, theta, slow gamma and fast gamma bands; n=54 seizures recorded in N=6 mice). **B**. Preictal representative cortical ECoG recording with the corresponding spectrogram and spectral decomposition, which showed features that were similar to those of hippocampal LFP recordings (mixed model repeated measures ANOVA, p<0.0001 for delta, theta, slow gamma and fast gamma bands; n=70 seizures in N=9 mice).

Similarly, spectral analysis of preictal EEG in patients revealed a progressive decrease of gamma oscillations and an increase of alpha oscillations in most of the electrodes (Fig.5), indicating that, as in DS mice, also in DS patients the preictal state shows specific electrographic signatures that precedes by minutes the seizure onset.

**Figure 5.**
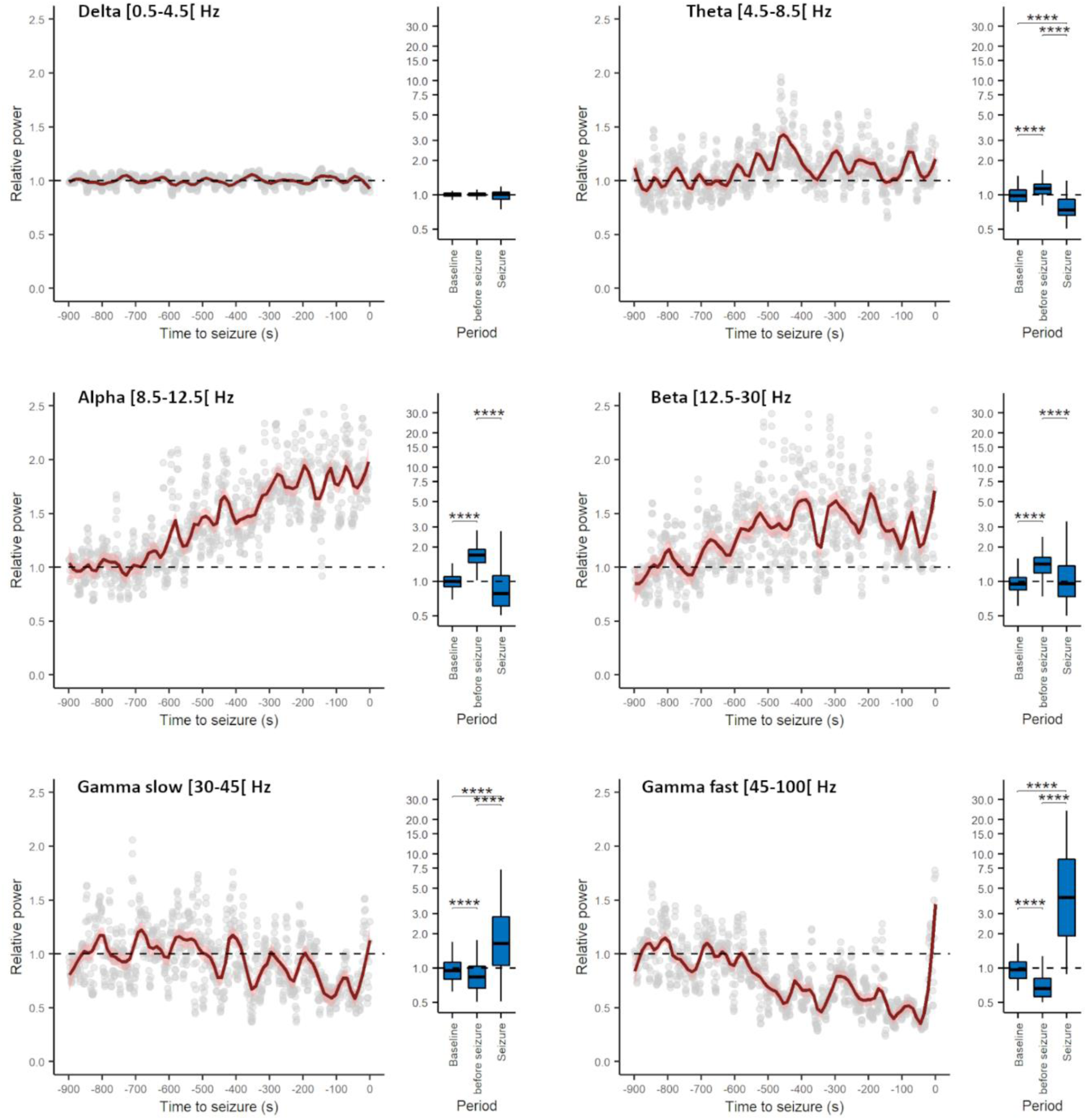
Generalized convulsive seizures are preceded by early changes in the spectral properties of all EEG channels in individuals with Dravet syndrome, as in mice LFP and ECoG. Evolution of the average relative power spectrum in the different EEG bands (number of channels= 46, number of seizure=3, number of individuals=3). The fast Fourier transform of the signal was computed and the power quantified from 900s before seizure onset to 200s after seizure onset over non overlapping 10s windows in delta (0.5-4.5 Hz), theta (4.5-8.5 Hz), alpha (8.5-12.5 Hz), beta (12.5-30 Hz), slow gamma (30-45 Hz) and fast gamma (45-100 Hz) bands. The spectral power of different frequency bands was normalized on the average between 900 and 700s before the seizures (considered interictal period). The time-series plots show a 900s window before seizure onset. The normalized power spectra between the interictal period, the 200s prior to seizure onset and the 200s after seizure onset (displayed in the box-chart plots) was compared using Kruskal-Wallis analysis of variance (ANOVA) on ranks followed by post-hoc Dunn’s test for pairwise multiple comparisons. *:p<0.05, **:p<0.01, ***, p<0.001, ****:p<0.0001.

Interestingly, the dynamics of the hippocampal LFP of DS mice and of the EEG of DS patients in the preictal period is coherent with previous evidence obtained in the DS mice, suggesting that seizures are preceded by alterations of network synchrony (36). We took advantage of the single-spike resolution of tetrodes recordings to analyze whether the perturbations of the network synchrony could be related to modification of the firing pattern of the two identified neuronal populations. More specifically, we fitted the ISI of recorded neurons to a gamma distribution (see methods for details) and we quantified during different preictal time windows the shape factor κ, which is a measure of firing regularity (34, 35). The analysis revealed a significant reduction of the shape specifically for the narrow spiking neurons (Fig.6 A-B, ANOVA mixed models F29,116=3.77 p<0.0001), whereas the firing dynamics of broadly spiking neurons was not modified (ANOVA mixed models F29,116=1.44 p=0.091). In particular, the ISI distribution of narrow spiking neurons revealed that they normally show a regular firing (shape factor κ>1), which progressively converges in the preictal period towards a Poisson-like or bursting regime (shape factor κ≤1).

**Figure 6.**
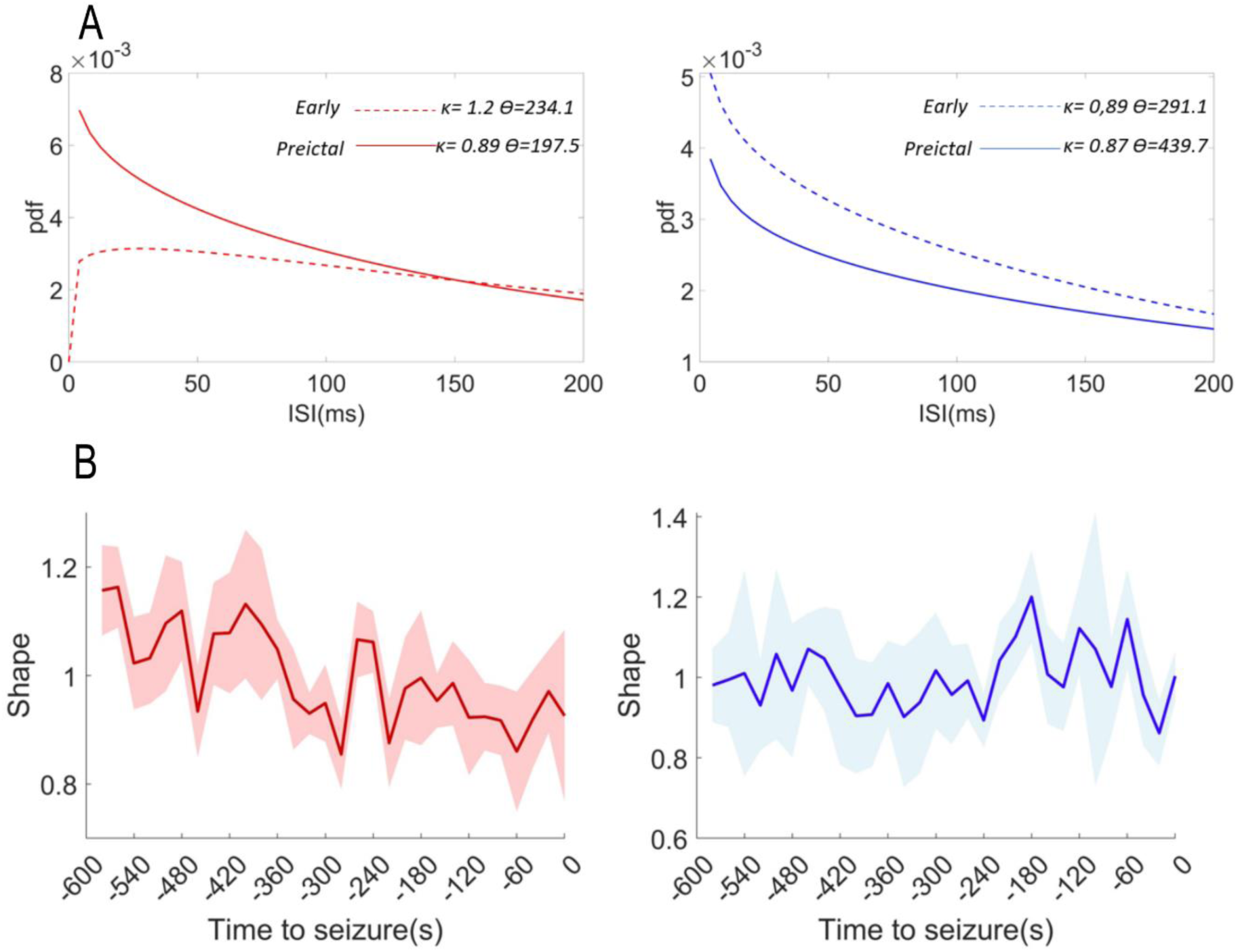
Seizures are preceded by early modifications in the firing dynamics of putative inhibitory neurons. **A**. Gamma distribution fitting of inter-spike intervals (ISI) in one-minute windows immediately before the spontaneous seizures (preictal) or 10 minutes before (early) for sharply spiking neurons (in red on the left) and widely spiking neurons (in blue on the left). The κ and ϴ values respectively represent the shape and the scale of the distributions(34, 35). **B**. Sharply spiking neurons (in red on the left) show a significant and progressive reduction of gamma distribution shape in the preictal period, whereas we did not find any modifications of the firing dynamics of widely spiking neurons (in blue on the left). The shaded areas represent the 95% confidence interval.

### Preictal and seizure onset properties do not depend on the specific mutation or on the natural history

Pathological dysfunctions of neuronal circuits in DEEs can be age dependent, as it has been previously proposed for DS (14, 36, 37), supporting the “multiple stages” concept of the pathology (6). To investigate if preictal or seizure onset features are mutation– or age-of-onset-dependent in our experimental framework, we exploited knock-in *Scn1a*^RH/+^ mice carrying the R1648H Na_V_1.1 loss-of-function mutation(38), in which the onset of a DS-like phenotype can be triggered by inducing a series of short seizures(17). We used a protocol of 5 short seizures induced with hyperthermia in a 5-day period (SIH protocol) either at P21 (age of spontaneous seizure onset in *Scn1a*^+/-^ DS mice) or in adulthood, and we chronically recorded the hippocampal LFP in both groups starting at P60 (Fig.7A). The SIH protocol was able to trigger long-lasting spontaneous seizures in both groups (Fig.7A), confirming and extending our previous results(17). The analysis of the hippocampal LFP showed that the features of the seizure’s onset and of the preictal period were the same in *Scn1a*^RH/+^ mice in which chronic epilepsy was triggered in adulthood compared to those in which it was triggered at P21 (Fig.7B, C & D): both *Scn1a*^RH/+^ groups recapitulated the features that we observed in *Scn1a*^+/-^ KO mice and DS patients. Moreover, as in *Scn1a*^+/-^ mice, we found similar features also analyzing the epidural ECoG recorded on the parietal cortex of *Scn1a*^RH/+^ mice (not shown), including preictal power increase for delta and theta frequency bands and decreased power for slow and fast gamma bands (mixed model repeated measures ANOVA; p<0.0001 for all the mentioned bands; 68 seizures in 6 mice). Thus, the features that we have observed do not depend on the natural history of the disease or on the specific mutation.

**Figure 7.**
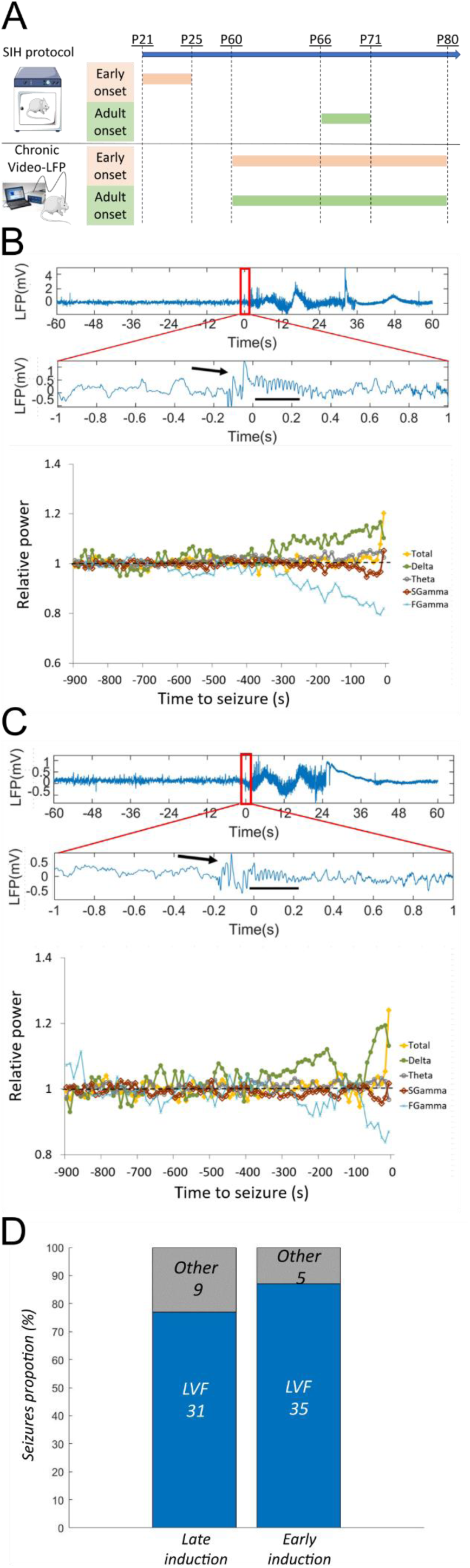
Properties of the seizure’s onset and of the preictal LFP do not depend on the specific mutation or on the natural history. **A**. Experimental protocol for triggering a DS-like phenotype in asymptomatic *Scn1a*^RH/+^ mice: we exposed *Scn1a*^RH/+^ mice to daily single short seizures induced by hyperthermia during a 5-day period (SIH protocol) either at P21 or in adulthood (P66), and we chronically recorded the hippocampal LFP in both groups starting at P60. The longitudinal recording of mice induced in adulthood allowed the quantification of seizures also before, during and immediately after the SIH protocol. This confirmed that spontaneous seizures are almost absent before the SIH protocol (only one mouse out of 7 displayed few spontaneous seizures), revealed that spontaneous seizures appeared in all mice between the third and the fifth day of the SIH protocol, and showed that they were still present even several weeks after the SIH protocol. These findings extend our published results(17), showing that an SIH protocol of 5 days is sufficient to trigger a severe phenotype and that the protocol is effective also in adulthood. On average, *Scn1a*^RH/+^ mice that underwent the SIH protocol at P21 displayed 0.4 spontaneous seizures/day in the period P71-P80, frequency that is similar to that of adult *Scn1a*^+/-^ mice (0.45 spontaneous seizures/day, ANOVA for genotype effect F(_1,18_)=1.62, p=0.24). Conversely, in the same recording period, we found a higher frequency of spontaneous seizures in *Scn1a*^RH/+^ mice that underwent the SIH protocol more recently (i.e. in adulthood) (0.61 seizures/day, ANOVA for age effect p=0.048). B. Upper and middle panels. Representative seizure with LVF onset (displayed with an enlarged time scale in the middle panel) recorded in *Scn1a*^RH/+^ mice that underwent the SIH protocol in adulthood. Lower panel. Quantification of LFP’s mean spectral power in the preictal period decomposed in different frequency bands, which showed a decrease of slow and fast gamma bands’ power and an increase of delta and theta bands’ power (mixed model repeated measures ANOVA; p<0.0001 for all the mentioned bands; 40 seizures recorded in 6 mice). C. same plots as in B, but for *Scn1a*^RH/+^ mice that underwent the SIH protocol at P21. Lower panel. Quantification of LFP’s mean spectral power in the preictal period showed, as for the other group of *Scn1a*^RH/+^ mice, a decrease of slow and fast gamma bands’ power and an increase of delta and theta bands’ power (mixed model repeated measures ANOVA; p<0.0001 for all the mentioned bands; 40 seizures recorded in 4 mice). D. Quantification of the proportion of LVF seizures in the two groups or *Scn1a*^RH/+^ mice.

## Discussion

Seizures with LVF onset pattern have been generally associated with focal seizures, and several studies proposed that they are caused by a sustained firing of GABAergic neurons at seizure onset, followed by an increase of the firing of pyramidal cells during the progression of the seizure (21, 23, 24). Here, we show that they are the principal seizures type also in DS, in a pathologic context characterized by reduced activity of GABAergic neurons. Strikingly, the analysis of single units’ dynamics in our model partially confirmed the GABAergic hyperactivity mechanism. In fact, we observed an increase in putative interneurons’ firing at the seizure onset, together with that of putative pyramidal cells, suggesting that this could underlie the LVF dynamics in this context.

Notably, we also disclosed temporal disorganization of the firing of putative interneurons in the preictal period that temporally correlated with modifications of hippocampal and cortical LFP spectral properties, preceding by minutes the onset of the electrographic seizure and that were observed also in all the channels of the EEG recorded in DS patients. These modifications are progressive in time, suggesting a continuous drift of the network toward the ictal state. These results have a double relevance: i) the detection of early EEG modifications could be useful to develop non-invasive methods for seizures prediction; ii) they are consistent with the existence of a specific functional state of extended networks in different brain regions in the preictal period (a “preictal brain state”)(39).

Importantly, seizures are currently unpredictable, have a strong impact on the quality of life of families and may worsen the DS phenotype. Thus, the identification of a potential EEG biomarker able to predict seizure several minutes before their onset could be clinically very useful, allowing families to organize themselves and possibly to use preventive therapeutic interventions that could inhibit seizures. Moreover, studying the properties of the preictal state could be informative about mechanisms that make brain networks prone to seizures. Indeed, we observed that on average the preictal LFP is characterized by higher power in the slow bands, especially in the delta band, and lower power in the gamma bands. The complex origin of the LFP signal makes it difficult to disclose a direct and specific link between its spectral composition and cellular activity. However previous studies highlighted a causal link between the activity of GABAergic interneurons and gamma oscillations(40). This is consistent with our results showing the preictal reduction of fast oscillations and the modification of the firing pattern of putative inhibitory neurons. This modification specifically concerns the temporal organization of the spiking activity, whereas the overall firing frequency was unaltered.

Tetrode recordings are not adapted to large-scale network analysis; however, our results are consistent with a previous study showing by *in vivo* Ca^2+^ imaging that the desynchronization of interneurons (reduction of pair-wise correlated activity) precedes hyperthermia-induced seizures in DS mice(41). We extend these observations by demonstrating that the firing pattern of interneurons is perturbed in the preictal period in DS mice also in spontaneous seizures. Moreover, by taking advantage of the temporal resolution of electrophysiological recordings, which overcome the imprecision of spike inference obtained from Ca^2+^ imaging, we demonstrate that the alteration of interneurons’ firing is not only modified at the network scale, but also at the single neuron level. More specifically, we observed in the preictal period a loss of firing regularity that specifically affects putative inhibitory interneurons, consistent with the hypo-function of GABAergic neurons in DS, followed by a rebound hyperexcitability of both inhibitory and excitatory neurons at seizure onset.

The fact that the hypo-excitability of GABAergic interneurons caused by Na_v_1.1 loss of function induces modifications of temporal spiking pattern, leaving unaffected the general firing rate, could appear counterintuitive. Several non-mutually exclusive hypotheses can be proposed. First, the complexity of the structure and dynamics of cortical networks can break the intuitive linearity between the excitability of a given neuronal population at the single cell level and its actual firing activity within a complex network. Indeed, previous studies have already demonstrated that reducing the excitability of inhibitory neurons by optogenetics can even cause a paradoxical increase in their firing rate within network activities, caused by the increase in excitatory drive from principal neurons (42, 43). These observations show that the organization of reciprocal feedbacks between inhibitory and excitatory neurons is extremely strong and efficient leading to a stabilization of the overall spiking activity of the network, a feature called Inhibitory Stabilized Networks (44). This feature may also explain why in DS mouse models the alteration of putative interneurons’ firing in the preictal period doesn’t induce evident modifications of the firing pattern of putative pyramidal cells. Another contributing mechanism may be the reported slower propagation of action potentials (which is increasingly slower for successive APs within a discharge) observed in GABAergic neurons of a loss-of-function Na_V_1.1 mouse model (45), which could disorganize the spiking pattern of the inhibitory neurons without affecting their overall firing frequency, especially when a sustained rhythmic inhibition is required by the network. Together with our results showing the presence of a preictal state characterized by changes in the LFP spectral composition, this suggests a scenario in which during specific states of the network the interneurons of DS mice and possibly patients would loss firing regularity causing a *qualitative* rather than *quantitative* alteration of the inhibitory activity in the network. This would progressively push the network toward an increasingly unstable but not hyperactive state until reaching a break-point at the seizure onset. According to this scenario, a primary reduction of network inhibition caused by alterations of firing regularity of inhibitory interneurons would result in an aberrant response of local feedbacks and in the paroxysmal increase of interneurons firing characterizing the onset of LVF seizures. This hypothesis is consistent with the recent observation that somatostatin-positive interneurons can be recruited early, even some seconds before hyperthermia-induced seizures in DS mice(46).

Moreover, analyzing the number of active neurons during a seizure, we found that the different putative neuronal subpopulations that we have observed are recruited in overall the same proportion. Interestingly, we observed the paroxysmal firing sustaining seizures activity only in a subgroup ranging from 30 to 50% of the recorded neurons. This is consistent with previous observations in models and patients(47, 48), and it supports the hypothesis that also in DS only specific subnetworks participate to seizures.

Notably, DS patients have both focal and generalized seizures, and many seizures have been reported as “falsely generalized”, because accurate analysis of video-EEG recordings performed during seizures that were classified as generalized showed that they have focal features at onset or develop in an asymmetric way (16, 49–51). The EEG correlate of some of these “falsely generalized” seizures has been described as bilateral abnormalities since onset, with a slow spike or a slow-wave, often followed by a brief attenuation, and then by fast activities in the theta range intermixed with slow waves (16). Although this pattern shows some similarities to a LVF onset, in our study we disclosed for several seizure types activities at higher frequency (in the gamma range), in particular in the attenuation phase, which are more characteristic of a LVF onset. Thus, this is consistent with focal LVF features for DS seizures. Our results hint three important observations: i) the mechanisms that sustain seizures at onset and after onset are not necessarily the same as those that are present in the preictal period and that drive the network toward the ictal state; ii) the properties of the seizure onset may not be informative about pre-ictal mechanisms; iii) specifically for DS, this feature is consistent with the lack of effect or worsening of some antiseizure medications that boost the GABAergic system(51).

Our study has some limitations. In particular: i) single unit recordings are not available for DS patients, because they do not undergo therapeutic brain surgery and associated pre-operatory intracranial recordings; ii) patients’ data are from a retrospective study with a small number of seizures recorded in routine EEG after the early stages of DS, when seizures are less frequent.

More studies are warranted for the quantification of LVF seizures’ frequency/preponderance and of detailed preictal features in patients.

Overall, the onset of generalized seizures in DS can have LVF features that are similar to those observed in other epilepsies for focal seizures triggered by hyperactivity of GABAergic neurons, but are characterized by perturbed activity of these neurons in the preictal period, which are linked to early pre-ictal modifications of EEG/LFP spectral properties that may be used as biomarkers for seizure prediction in DS patients.

## Methods

### Mice

Heterozygous conditional global *Scn1a* knock-out (*Scn1a*^+/-^) mice were generated by crossing the *Scn1a* exon-25 floxed mouse line with mice expressing Cre recombinase under the *Meox2* promoter(52), both in the C57Bl/6j genetic background. This allowed to induce early embryonic *Scn1a* haploinsufficency and to reduce mortality in the C57Bl/6j colony(52). Scn1a^RH/+^ heterozygous knock-in male mice in the C57BL/6J background were crossed with 129/SV females (Charles River, France); the F1 generation of the hybrid mixed (129:B6 50%-50%) background was used for all experimental groups(17). Genotyping was performed as already described(17, 52). We followed ARRIVE guidelines for planning experiments and reporting results. Experimenters were not blinded to the genotype; animals were randomized (www.randomizer.org) within each experimental group. Animals were kept in standard housing conditions with water and food ad-libitum until the surgery for electrode implantation. After the surgery, mice were kept separated to prevent the risk of reciprocal damage of the recording system. To reduce stress due to social isolation during chronic recordings, mice were kept in couple of the same sex in two hemi-cages, separated by a pierced metallic separator allowing for sniffing and tactile interaction. Experiments were performed in accordance with national and European legislation, approved by ethical committees and by the French ministry (MESRI, APAFIS #13619 and #15665-201805301624157_v2).

### LFP and ECoG recordings

Hippocampal LFP was chronically recorded with polyimide insulated stainless-steel electrodes (125µm diameter, P1 Technologies, US) implanted in the dorsal hippocampus by stereotaxic surgery under 2% isoflurane anesthesia at the following coordinates: 1.9mm posterior to bregma, 1.5mm lateral from bregma and 1.5mm ventral from the dura. An electrode fixed with a screw on the skull in the occipital region was used as ground. ECoG signals were acquired in a separate group of mice with a stainless-steel screw implanted in the dura above the parietal cortex (3mm posterior and 3mm lateral from the bregma). Both types of electrodes were connected to a plastic pedestal (P1 Technologies, US) and fixed on the skull with dental cement allowing for stable and long continuous recording. After a recovering period from surgery (at least 1 week), mice were connected to the amplifier and acquisition system (ADInstruments, UK). Infrared video cameras were synchronized to the acquisition of the ECoG signal allowing the simultaneous recording of electrographic and behavioral seizures. Mice were recorded for 3 weeks.

### Single unit recordings

We recorded single units with tetrodes prepared by twisting together four 17µm platinum/iridium wires (California Fine Wire, US). Four tetrodes were inserted inside a guide cannula, wired to an 18-channels connector (Omnetics, US) and fixed on a microdrive (Axona, UK). The ground electrode was connected to the guide cannula. They were implanted by stereotaxic surgery under 2% isoflurane anesthesia at the following coordinates: 1.9mm posterior to bregma, 1.5mm lateral from bregma and 1.2mm ventral from the dura and fixed on the skull with dental cement. After one week of recovery from the surgery, mice were daily recorded (DACQUSB recording system, Axona, UK). Tetrodes were regularly lowered by 50µm steps. Mice were put in a rectangular open field (70X45cm) with sugar pellets randomly scattered on the floor, and left to freely move. Head position and orientation were tracked by two infrared leds fixed on the headstage. The freely moving session was used to evaluate the quality of the LFP signal and of the unitary activity in order to select the mice to record during the hyperthermia-induced seizures and the chronic recordings.

### Seizures induction by hyperthermia

Mice were placed in a small incubator and the core temperature, continuously measured using a rectal probe, was increased 0.5 °C every 2 min as described previously(17). They were immediately removed from the incubator when a behavioral seizure occurred or a maximum temperature of 43°C was reached. Knock-in Scn1a^RH+/-^ mice were exposed to five consecutive daily hyperthermia-induced seizures, which led to epileptogenesis and spontaneous seizures as described previously(17). In knock-out *Scn1a*^+/-^ mice, seizures were induced with hyperthermia only for single units recording experiment. In this case, we did not expose the same mouse to hyperthermia more than twice per week, to reduce the risk of death during seizures.

### Patients

We investigated the seizure onset features of individuals with DS analyzing the EEG recordings obtained from the database of Necker-Enfants Malades hospital, a tertiary center for rare epilepsy. Our study is based on retrospective EEGs performed during the classical follow-up of individuals with DS. We included DS patients with a *SCN1A* pathogenic variant and seizures recorded after the stormy phase. EEG recordings used in this study lasted from 40 minutes to 120 minutes. The 11-channel EEG data were recorded with a digital acquisition system (Natus Medical Inc), according to the 10/20 international system with Fpz as reference and with a sampling frequency of 256Hz. In compliance with French law, the consent regarding non-opposition for this study and the use of their data was obtain for each individual and their parents. The ethics committee of Necker-Enfants Malades hospital approved the study protocol; the subjects’ consent was obtained according to the Declaration of Helsinki.

### Data Analysis

#### LFP/ECoG

Chronic LFP and ECoG were sampled at 2KHz and bandpass filtered 0.1-1000Hz. LFP during hyperthermia was sampled at 4.8KHz, bandpass filtered at 0.1-1000Hz and notch filtered at 50Hz. Seizures were detected by offline visual inspection of the traces and of the synchronized video with LabChart8 software (ADInstruments, UK). Seizures onset was defined as a change in the signal accompanied by subsequent typical seizure activity, clearly distinguished from background and interictal activity(29). To determine and classify the seizure’s onset, two seconds of the signal around the onset were visually inspected and classified according to previous clinical and experimental criteria(28, 53). All subsequent analysis was performed using custom-built scripts in MATLAB-2018 (MathWorks, US). LFP/ECoG spectrum in the preictal period was obtained by fast Fourier transform of the signal in the 15minutes preceding the seizures over non overlapping 10s windows. This strategy allowed the elimination of windows containing big artifacts (amplitude of the signal >2mV) due to movement, scratching, or eating/drinking. These windows were left blank and non-considered for further analysis. The total power of the signal and the relative power of different frequency bands for each preictal period was normalized on the average between 15 and 10 minutes before the seizures.

#### Single Units

Signals were sampled at 24KHz and band-pass filtered 300-7000Hz. Spikes were identified by amplitude thresholding and the waveforms stored as 50 points (0.2ms before and 1.8ms after the peak). Spike sorting was performed manually using the graphical cluster-cutting software Tint (Axona, UK). To analyze the firing regularity in the preictal period, we divided the preictal period into 30 windows of 20 seconds. We fitted the gamma distribution to the ISIs by using the *fitdist* MATLAB function and we used the shape factor of the gamma distribution as an index of firing regularity (35). In particular, the shape factor is expected to take a value of 1 for a random process following a Poisson distribution, a value>1 for a more regular firing distribution or a value<1 for a bursting firing. To ensure a sufficient number of ISI for a reliable distribution fitting, we only considered neurons with a minimal number of 20 ISI in each window, similar to previous published studies (35).

#### Clinical EEG

To quantify preictal and seizure’s onset spectral bands compared to baseline, we computed the fast Fourier transform and quantified the power of the different frequency bands.

#### Statistics

Data were compared using parametric or non-parametric tests according to their distribution. Specific tests are indicated in the text and in the figure legends.

## Data availability

Data are available upon request.

## Acknowledgements.

We thank Emilie Bonnet (Institute of Molecular and Cellular Pharmacology, CNRS UMR 7275 and University Cote d’Azur) for her skillful technical support and Dr. Anna Kaminska, head of the Clinical Electrophysiology unit of Necker Enfants Malades hospital, for providing the video-EEG recordings of DS patients.

## Author Contributions

FC, MM conception and design of the study.

FC, MK, JL acquisition and analysis of data.

FC, MK, RN, MM interpretation of results.

FC, MK, RN, MM drafting a significant portion of the manuscript or figures.

